# Improved protein docking by predicted interface residues

**DOI:** 10.1101/2021.08.25.457642

**Authors:** Gabriele Pozzati, Petras Kundrotas, Arne Elofsson

## Abstract

Scoring docking solutions is a difficult task, and many methods have been developed for this purpose. In docking, only a handful of the hundreds of thousands of models generated by docking algorithms are acceptable, causing difficulties when developing scoring functions. Today’s best scoring functions can significantly increase the number of top-ranked models but still fails for most targets. Here, we examine the possibility of utilising predicted residues on a protein-protein interface to score docking models generated during the scan stage of a docking algorithm. Many methods have been developed to infer the portions of a protein surface that interact with another protein, but most have not been benchmarked using docking algorithms. Different interface prediction methods are systematically tested for scoring >300.000 low-resolution rigid-body template free docking decoys. Overall we find that BIPSPI is the best method to identify interface amino acids and score docking solutions. Further, using BIPSPI provides better docking results than state of the art scoring functions, with >12% of first ranked docking models being acceptable. Additional experiments indicated precision as a high-importance metric when estimating interface prediction quality, focusing on docking constraints production. We also discussed several limitations for the adoption of interface predictions as constraints in a docking protocol.

## INTRODUCTION

Most proteins carry out their biological functions through interactions with other proteins ^1^. Subsequently, the ability to modulate protein-protein interactions (PPI) could lead, among other things, to the cure of diseases. However, modulating PPIs requires a fundamental understanding of PPI details on the atomic level. Experimental methods, like X-ray crystallography or NMR/ EM spectroscopy, can produce highly reliable structures, but unfortunately, these methods are expensive and time-consuming^2^.

A completely different approach to derive such structures involves the use of computational methods ^3^. Unfortunately, this approach is limited by the dynamic nature of protein behaviour in vivo. For instance, most proteins undergo structural rearrangements or conformational changes when interacting with a partner ^4^. Also, in some cases, PPI is obligate, meaning that the protein must fold into a stable and functional conformation ^4,5^. Other PPIs are non-obligate, meaning that interaction partners may also exist in a stable but non-associated form. Obligated complexes are generally permanent, but most non-obligate complexes are transient. Their lifetime is influenced by several factors, including physiological conditions (pH, salt concentration etc.), the concentration of interaction partners and the state of certain molecular switches ^5^. Furthermore, obligate and non-obligate complexes have different geometrical and physicochemical properties of their interfaces ^6^. Thus, the prediction of 3D structures of protein-protein complexes (protein docking) remains one of the most demanding challenges in computational biology.

Usually, a structure of uncharacterised PPI is derived from structures (experimental or modelled) of individual proteins by rigid-body ^7^ or flexible docking procedures ^8,9^. These protocols generally consist of two stages: fast generation of large numbers of putative mutual arrangements of two proteins (docking model or pose) using simplified energy function (scan stage) and subsequent application of a more complex scoring function to the obtained configurations, in order to discriminate the few ones that most likely are close to the native structure (scoring stage) ^10^. Rigid-body docking is generally faster than flexible docking, but flexible docking (that allows intra-protein conformational degrees of freedom) better reflects the dynamic nature of the proteins ^9^. Limitations of these methods are implicit in the necessity to generate large amounts of the docking models (usually on the order of hundreds of thousands) to have a significant chance of generating at least one near-native docking model. Many decoys are not a problem but necessitate an extraordinarily accurate and computationally efficient method to identify the few near-native solutions. Some methods also use much smaller datasets for testing ^11^, i.e. these methods do not work for the general docking problem. Another common strategy is reducing the number of considered docking poses by performing clustering and only applying a scoring function to the cluster representatives ^10^. With such an approach, acceptable docking models can be found in the top 10 scored poses for almost 40% of complexes in the widely adopted Benchmark 5.0 dataset ^12^,^13^.

Another approach is to use constraints derived from, e.g. prediction of protein interfaces ^14^. The goal of interface prediction is to understand which residues from the surface of one protein are more likely to form contacts with the residues of an interacting partner (interaction patch). This kind of prediction is commonly based on evolutionary features acquired from standard multiple sequence alignments (MSA) ^15,16^. Most predictors use different combinations of such information with sequence and structural features of proteins alone (unbound) and in the associated (bound) form ^17–19^. Unfortunately, many interface predictors have been published without testing how they would improve the success of protein docking algorithms. ^17,19–25^. The possibility to use standard MSA is quite important, given that combining MSAs from different interacting proteins (which is required for some interface contact prediction algorithms) is a non-trivial task ^26^. Another main advantage of predicting interface patches is that, considering proteins singularly equalise on a similar order of magnitudes, the number of interacting and non-interacting residues, making the two categories more or less balanced, according to the protein type. This last property is important for all the machine learning methods commonly applied to this problem, particularly Support Vector Machines (SVM) and Artificial Neural Networks (ANNs). Indeed, most machine learning algorithms are consistently influenced by unbalanced datasets and tend to learn undesired patterns, such as proportions of classes, from the provided training sets ^27^.

Dockrank is one of the most recent attempts to use interface predictions in protein-protein docking ^28^. This work has shown some consistent improvement in the docking success when applying interface predictions to the scoring of the docking poses. However, the dataset used in that study was limited to complexes with sufficient confidence of predicted interface residues, which reduces the generalisation of the conclusions. Furthermore, other studies were conducted on small or bound datasets only, and in some cases, the predicted interface information was used in combination with other scoring parameters, which made the exact contribution of interface predictions unclear ^29–31^. Thus, it is still unclear how much valuable information for docking can be extracted from interface prediction. In order to clarify this point, we filter docking poses produced by the GRAMM docking software ^32^, utilising interface information acquired from native structures of PPI in the DOCKGROUND dataset and various interface predictors. This protocol aims to establish a reference framework for easy quantification of the performance of different interface predictors when applying them in a real-case docking scenario when the native PPI structure is not known.

## MATERIALS AND METHODS

### Dataset

This study utilised all dimeric protein complexes extracted from the benchmark set 4 ^33^ from the Unbound section in the DOCKGROUND website: http://dockground.bioinformatics.ku.edu/. Additionally, we excluded all the complexes containing chains shorter than 50 residues, leading to a set of 220 protein pairs for which both single-chain (unbound) and associated (bound) experimental structures are available. Depending on root-mean-square deviation (RMSD) between interface Ca atoms in unbound and bound structures (*i*-RMSD) and fraction of non-native contacts (fnon-nat) in unbound structures ^12^, this dataset can be divided into 116 easy (*i*-RMSD < 1.5 Å and fnon-nat < 0.4), 72 medium-difficulty (1.5 < *i*-RMSD < 2.2 and fnon-nat > 0.4) and 32 hard (i-RMSD > 2.2) cases.

The numbering of residues in the unbound structures has been mapped to the numbering in the bound structures using pairwise global sequence alignment utility from the biopython package (version: 1.76) with the BLOSUM62 scoring matrix ^34^. In order to facilitate the following comparisons, all residues from bound structures with no correspondence in the unbound structures have been trimmed. Furthermore, unbound chains have been structurally aligned to the bound counterpart to determine a level of difficulty for the docking of each complex. Here we adopted three difficulty classes; hard, medium, and easy, as described previously^35^. Finally, for each complex, the longer (shorter) chain has been re-labelled ‘A’ (‘B’) and henceforth is referred to as receptor (ligand).

### Rigid-body docking protocol

Unbound structures of the proteins in the dataset were docked utilising Fast Fourier transform (FFT) rigid-body docking algorithm as implemented in the scan stage of the GRAMM software ^32^. Unlike other FFT-based programs (e.g., ZDOCK ^7^ and ClusPro ^36^), GRAMM does not include any other energy components (electrostatics, desolvation, etc.) besides simplified Lennard-Jones potential when generating an initial set of docking poses. Therefore, using all these models allows investigating the ‘pure’ effect of various factors on a minimally biased set of docking models generated with only the surface geometry of the receptor and ligand taken into account. Further, the unique low-resolution nature of the GRAMM docking algorithm permits small amounts of atomic clashes on the interfaces of the docking models, which to a certain degree accounts for the conformational flexibility upon protein binding ^32^.

Default grid sizes (32×32×32 or 64×64×64) and calculation parameters (grid step 3.5 Å, rotation angle 10 degrees) have been used for all complexes except 4YOC, where it was necessary to increase grid size to 128×128×128. 340,000 docking poses were generated for each docking pair to ensure that at least one near-native docking model is presented for all the complexes considered. GRAMM output (translation vector and three Euler angles per docking pose) were transformed into Cartesian coordinates of the ligand using a script written with the Tensorflow python library (version: 1.13.2). In both steps, different dockings may be elaborated in parallel, consistently reducing the computation time. The initial docking poses were further re-scored by a function:

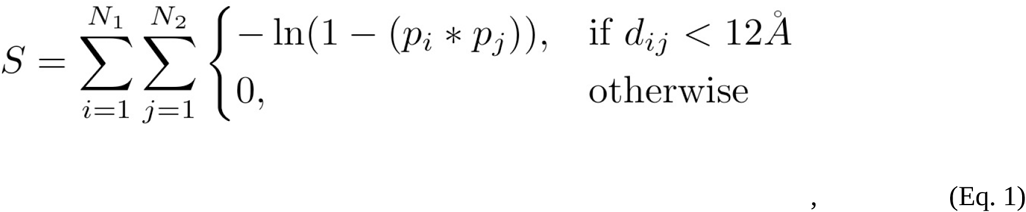

where the summation is performed over all *N*_1_ and *N*_2_ residues of the receptor and ligand, respectively, *p*_*i*_ and *p*_*j*_ are, correspondingly, the probabilities (given by an interface predictor) of residues *i* of the receptor and *j* of the ligand to occur on the native interface, and *d*_*ij*_ is the distance between C_β_ atoms of residue *i* in the receptor and residue *j* in the ligand. In order to avoid singularities in Eq. 1, an upper limit of 0.99 for *p*_i_ and *p*_*j*_ was used. Ten highly-scoring docking poses were retained for further evaluation. For comparison, we also used docking poses re-scored by the atom-atom contact energy AACE18 ^37^.

### Interface predictions

We selected several predictors (Table 1) for calculating propensities of the residues to occur inside the native interaction patch. We are aware that there are many more interface predictors described in the literature, but our choice was restricted by the availability and portability of the code to run locally. BIPSPI ^22^ produces estimates of interface patches and inter-protein contacts for a pair of either sequences or structures. In this study, pairs of structures were provided as input, and the two interfaces returned from the predictor were used for scoring. ISPRED4 ^18^ first uses a Support Vector Machine (SVM) to generate initial interface residue propensities. In ISPRED4, these predictions are further processed by Conditional Random Fields (CRF). However, no improvement was seen in our study using the second set and, therefore, the CRF predictions were ignored.

**Table 1:** interface residue predictors

| Predictor | Description | Ref. |
| --- | --- | --- |
| SPPIDER | Neural Network consensus method based on protein structure geometric features and predictions of relative solvent accessibility. | 40 |
| PredUs | Support Vector Machine method based on solvent accessibility and position conservation derived from protein structural alignment. | 38 |
| dynJET | A model which combines evolutionary, geometric, physicochemical, and interface propensity features. | 39 |
| ISPRED4 | A method based on Support Vector Machine and Conditional Random Fields, which combines information about residue structural context, physicochemical, and multiple sequence alignment features. | 18 |
| BIPSPI | Tree classifier trained with XGBoost algorithm, based on structural and multiple sequence alignment features obtained for pairs of proteins. | 22 |

Further, SVM-based binary interface predictions have also been obtained using the PredUS predictor ^38^. The dynJET2 algorithm ^39^ has been applied to our dataset in the “SCnotLig” mode to exclude possible ligand binding pockets and using three iterations to increase reliability. Results of both trace and cluster calculations have been tested in the final docking as interface predictions. For cluster-based predictions, only values supported by all three clustering iterations was considered (as suggested by the authors), setting the probability to zero otherwise. Finally, the SPPIDERII algorithm from the SPPIDER Web server [35] was used to generate predictions in the regression form, obtaining continuous probabilities from 0 to 1 (all other options have been left at their default values).

### Native interfaces

Native interface residues were extracted from the bound structures using the condition that solvent accessible surface area (SASA) of a residue in a protein in isolation should be larger than when the protein is bound to the interacting partner. SASA was calculated employing the DSSP v.3.0.0 module ^41^ implemented in the biopython library. If a residue from the unbound structure had no correspondence to the bound one, the same criteria were evaluated on unbound structures superimposed on the corresponding bound.

### Assessment of interface predictors

Interface prediction quality has been evaluated using two classic metrics: True Positive Rate or Recall, TPR:

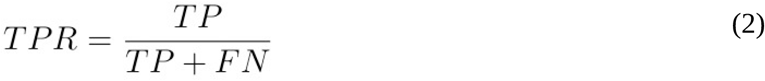

and Precision, PPV:

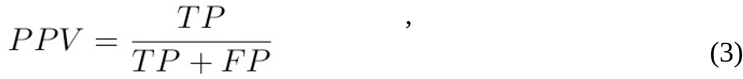

where TP, FP, and FN are numbers of true positives (correctly predicted interface residues), false positives (non-interface residues incorrectly predicted as interface) and false negatives (interface residues incorrectly predicted as non-interface) for a specific protein chain, respectively. For the interface predictors that output continuous probabilities rather than binary classification (interface/non-interface), all those quantities are dependent on the probability threshold, above which a residue is considered to be on the interface. Thus, to evaluate the overall performance of such predictors, we used the area under the precision-recall curve (AUC) computed for decreasing thresholds using the scikit-learn python package (v. 0.24.1). In our pipeline, an interface predictor produces two predictions for each protein complex considered (one for receptor and another for a ligand) with generally different AUC. We use both sets per complex or a set with the smaller AUC (henceforth referred to as *worst chain* predictions) for further analysis. For evaluating the overall performance of an interface predictor, we averaged TPR and PPV values for all protein chains in the dataset and analysed the distribution of AUC values.

### Assessment of docking predictions

To assess the quality of a docking model, we adopted the DockQ score ^42^, which combines all evaluation criteria used in the CAPRI competition ^43^ into a single score, into a range from 0 to 1, with 1 representing a perfect match between a docking model and the native complex structure. Here, dockQ values of 0.23, 0.49, and 0.8 represent threshold values ^42^ for docking models of acceptable, medium and high quality in terms of the CAPRI criteria. DockQ scores were obtained by comparing a docking model with the bound version of the complex structure if not specified differently. To measure the overall performance of a docking protocol over the entire dataset, we evaluated the fraction of acceptable models (defined by DockQ>0.23), *SR*(*N)*, in the top *N* ranked models. Here, we analysed *SR*(*N*) for all *N* ≤ 10.

### Simulated interface predictions

In order to observe the behaviour of interface prediction-driven docking in a controlled scenario, simulated interface predictions have been generated by introducing pre-defined levels of noise in the native interfaces. First, randomly selected interface residues from each protein chain were marked as non-interface to reach a certain TPR. After that, randomly selected surface residues not belonging to the interface were marked as interface until reaching a certain value of PPV. In this study, we considered nine different datasets with various (TPR/PPV) values: (0.25/1), (0.5/1), (0.75/1), (1/0.25), (1/0.5) and (1/0.75).

### Availability

All code is available from git.. All data for all methods is available from fighshare

## RESULTS AND DISCUSSIONS

### Baselines for the docking performance

The lower baseline for our docking pipeline was determined by analysing “raw” GRAMM output (ranked by shape complementarity only). Then, the docking protocol yielded at least one acceptable docking model among the top 10 models for 12 complexes (*SR*(10) ~ 5%) with an average DockQ score of 0.04. The upper baseline was estimated using all native interface residues by setting *p*_*i*_ and *p*_*j*_ in Eq. 1 to a probability of 0.99. In this case, SR(10) jumps to 81%, with an average DockQ score of 0.45. Top ranking models are of acceptable or better quality for almost half of the targets, SR(1) = 46% with average dockQ ~ 0.27. Easy cases from the dataset yielded SC(1) ~ 62%, but even medium and hard cases displayed significant SC(1), with 35% and 16%, respectively (Fig 1, right panel).

**Figure 1.**
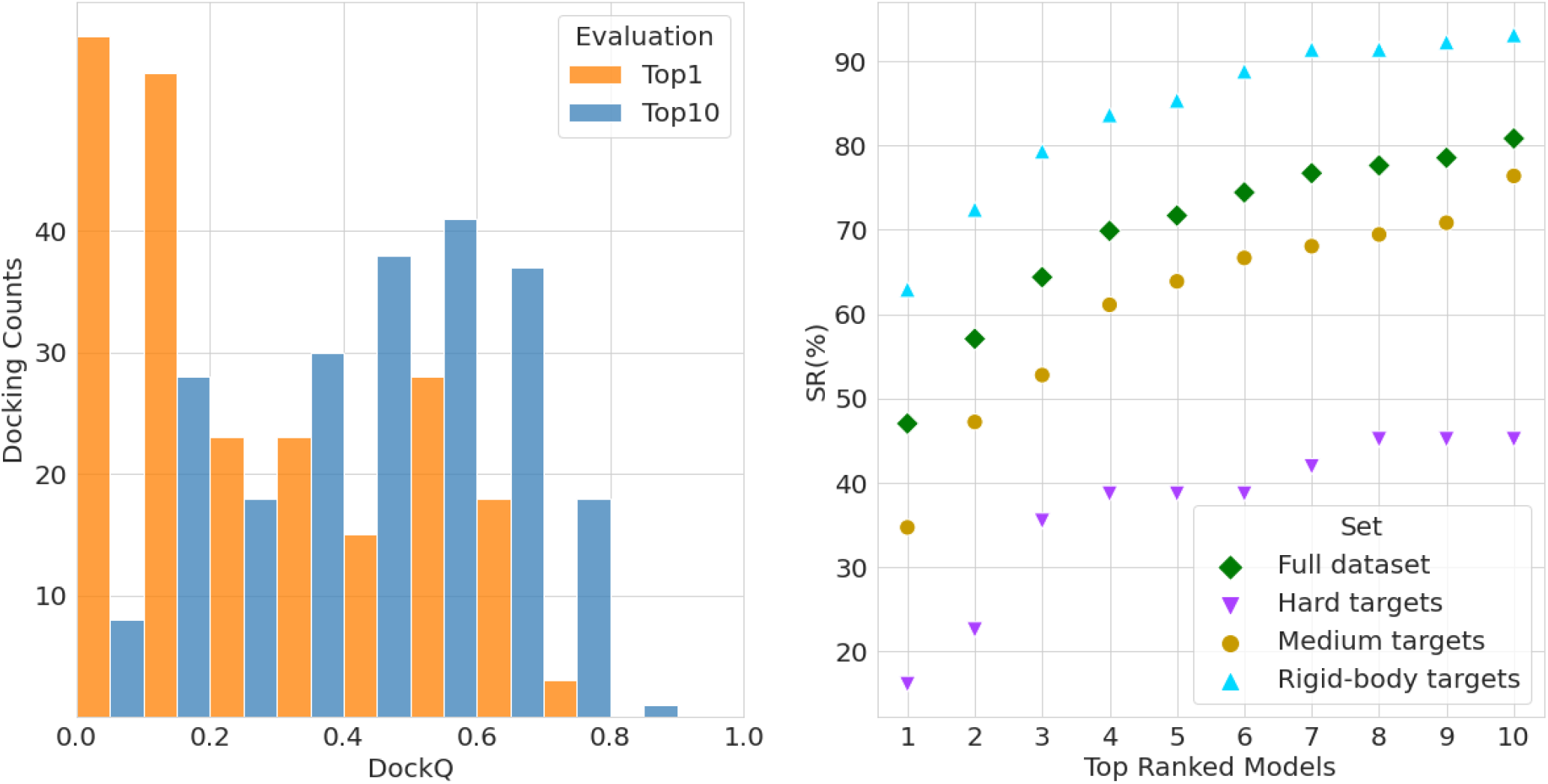
Performance of the docking with constraints derived from the native interfaces; (left) distribution of DockQ scores; (right) success rate, *SC*(*N*), as a function of the number, *N*, of considered top docking models. Data in the left panel pertains to the entire dataset of 220 binary complexes from DOCKGROUND benchmark set 4, while the right panel displays results for the entire dataset and the three sub-groups separately.

Among the forty-two targets with no acceptable docking models in the top 10 models, there are seven easy, eighteen medium-difficulty, and seventeen hard examples (6%, 25% and 53% of corresponding cases in the entire dataset). The lower performance on the hard targets indicates the significance of accounting for the flexibility in the docking protocol. Nevertheless, near-native docking models are present further down the list for all complexes in the dataset.

However, the difference between bound and unbound conformations of the proteins in the dataset led in several cases to the imperfect shape complementarity in the unbound “native” PPI structure (unbound structures superimposed on the bound ones in their native arrangement) while scoring equation (Eq. 1) favorises docking conformations with more contacts. In addition, docking constraints utilised in this study are considered on the residue level rather than on the residue contact level. Hence, the current re-scoring scheme may bring to the top of the prediction list docking models that have interface patches of the receptor and ligand surfaces correctly facing each other, but with the ligand rotated so that this mutual ligand and receptor position maximises the number of contacts for the unbound structures (an example is shown in Fig 2). Indeed, there is a significant number of top 1 docking models with a slight deviation of their interface center of mass (CM) from the CM of the native interface (Fig 3A). Notably, for the best out of the top 10 docking models, this number is significantly smaller, and the DockQ score exhibits the expected correlation with the CM deviation (Fig 3B), indicating that given correct interface constraints, it is desirable to analyse top 10 models in order to infer docking models with correct mutual orientation of the receptor and ligand.

**Figure 2.**
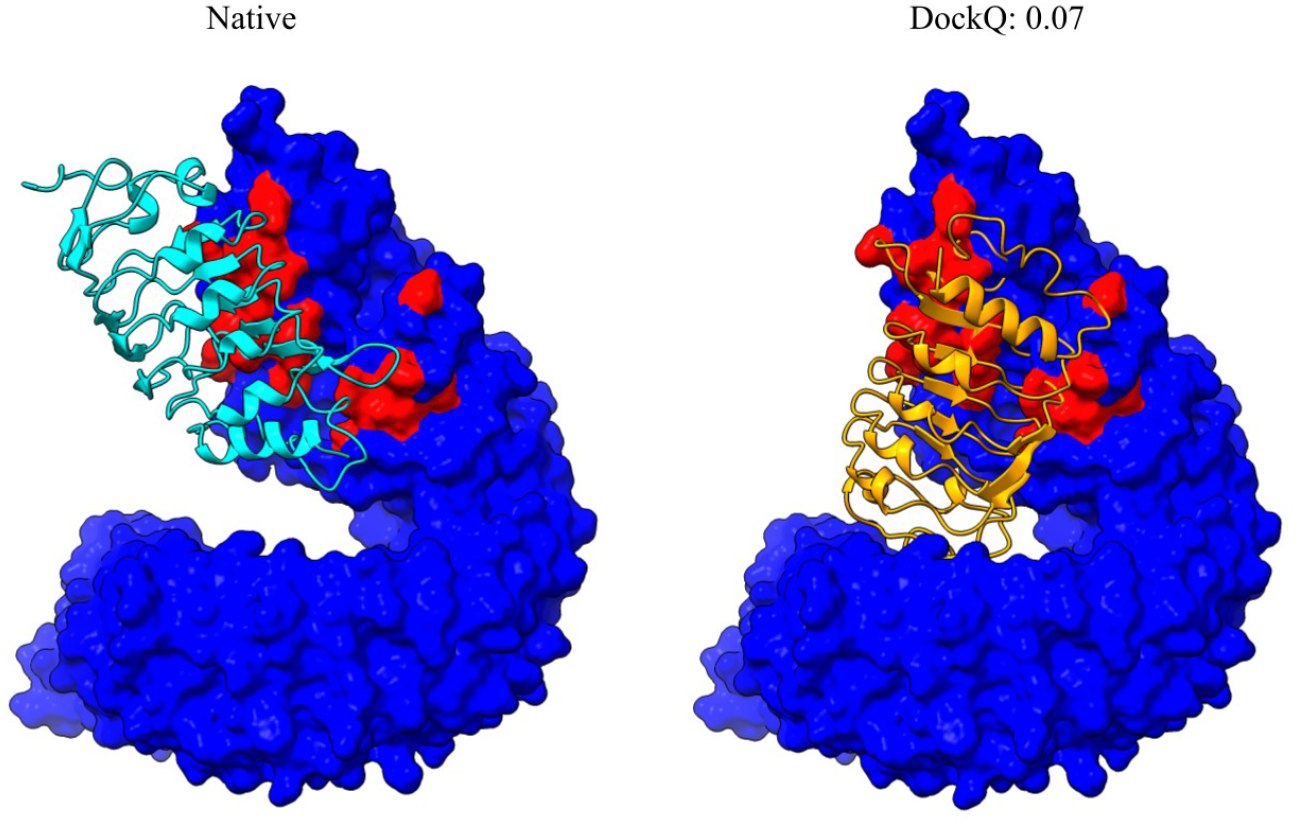
Example of the docking impaired by maximisation of the number of contacts. The left panel displays a reference “native” complex built by the superimposition of the unbound structures taken from PDB 4LSA, chain A (receptor) and 4LSC, chain A (ligand) onto, correspondingly, the chains A and C of the PDB 4LSX. The right panel depicts the best docking model among the top 10 models re-scored by equation (Eq. 1) with the 99% probabilities for the native interface residues. In both panels, receptors are represented by the atomic surfaces and coloured red (blue) for the native interface (non-interface) residues, while ligands are displayed as the cartoons.

**Figure 3.**
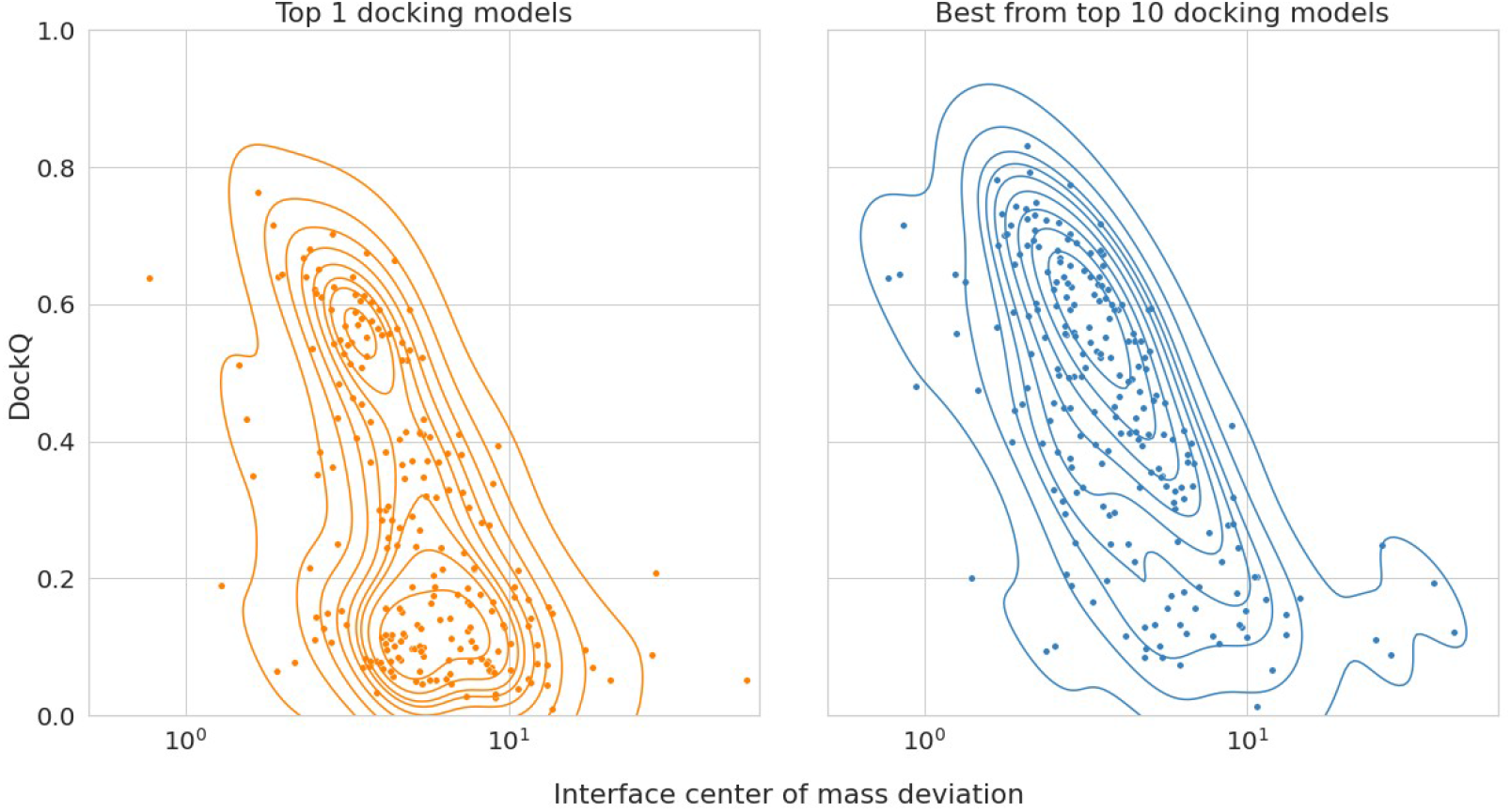
Interface center of mass deviation correlated to DockQ score from dockings with real interface constraints. For each complex in the DOCKGROUND benchmark-4 dataset, the center of mass coordinates computed for the native complex interface residues and the identical residues in the best docking solution (in top rank and top 10 ranks) were obtained using real interface constraints. The distance between the two centers of mass has been plotted against the relative docking model DockQ score. Kernel density estimator (seaborn library, default settings) has been adopted to visualise the density better.

### Performance of interface predictors

The best overall identification of interface residues is observed for the BIPSPI predictor, with an AUC of 0.49 (Fig 4, left panel), clearly superior to the other methods (AUC: 0.27-0.33). Further, predictions from PredUS have been evaluated using a single combination of TPR and PPV due to the binary output. This predictor reached a performance comparable to SPPIDER and ISPRED4, with TPR=0.37 and PPV=0.32. Examining the overall distribution of individual chains, all predictors, except BIPSPI, have similar median values ranging between 0.25 and 0.3, with 73% being better predicted than random. When the performance of interface predictors are assessed using *worst chain* predictions (see Methods), the precision-recall curves obtained a behaviour very similar to what was expected from a random predictor (AUC=0.22), data not shown. Again, the only exception is BIPSPI, which yielded an average AUC of 0.32.

**Figure 4.**
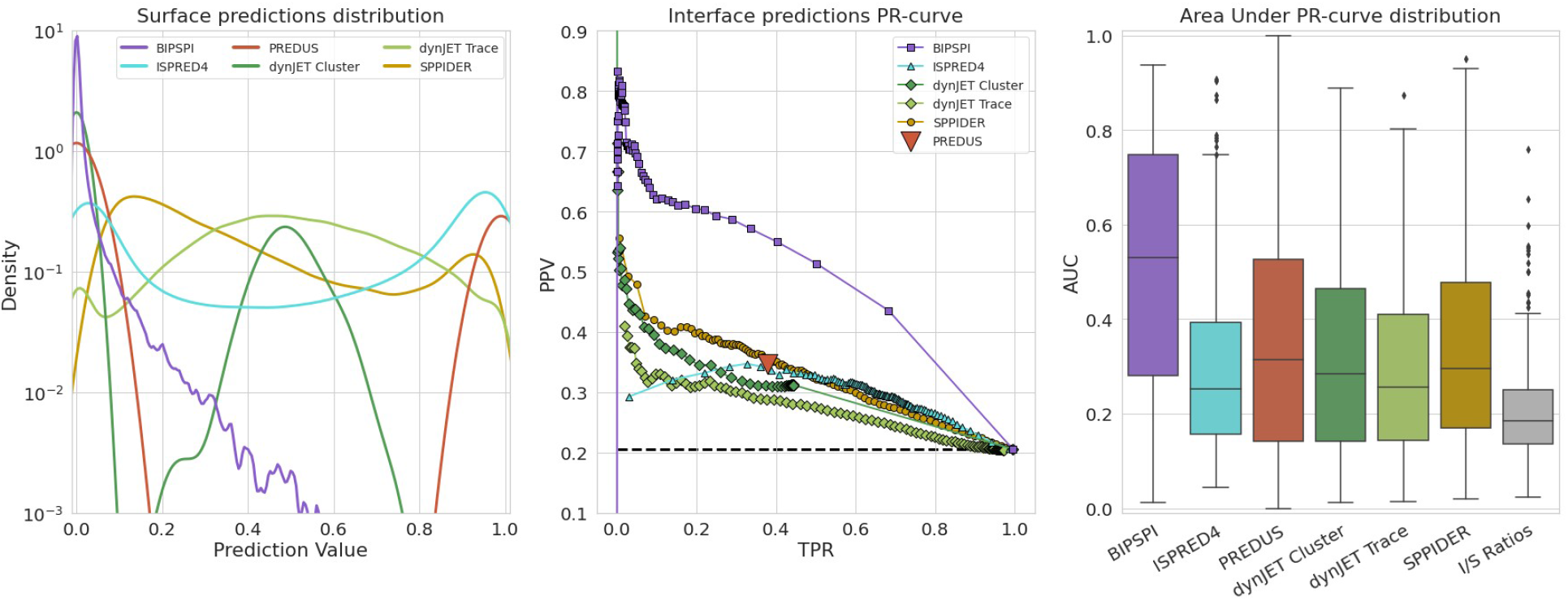
Overall behaviour of different interface predictors on 220 binary complexes from DOCKGROUND benchmark set 4. In the left panel, distribution of predictions for each surface amino acid in the dataset have been reported for each predictor. The mid panel displays precision-recall curves with TPR and PPV averaged over all protein chains in the dataset. The dashed line represents the average ratio of interface and surface residues (I/S ratio), which serves as the expected performance of a random predictor. For PREDUS no AUC curve can be drawn; thus, a single triangle represents its performance. In the right panel, AUC distributions for individual protein chains are shown as box-and-whiskers plots, each corresponding to a different interface predictor. For the PREDUS, PPV values are reported instead due to their equivalence to the AUC. As a random predictor reference, the distribution of the I/S ratios for individual protein chains is shown.

Notably, BIPSPI is the only predictor that considers pairs of structures simultaneously to infer their interface. All the other predictors use only a single structure. Therefore they might predict alternative interfaces, interacting with different interaction partners, possibly explaining the superior performances of BIPSPI. Further, all predictors, except BIPSPI ^22^, consistently perform worse than reported in the original publications. The decreased performance could be related to overtraining of the methods. One indirect confirmation for this hypothesis is given by the structural similarity of the complexes, which are responsible for the spike in AUC at ~20% of surface residues at the complex interface (Fig S1, right panel, I/S ratio 0.2), to the complexes from the original BIPSPI training set (Benchmark 5 ^12^). The average TM-score for this set is 0.89. In comparison, the complexes responsible for the drop in AUC at I/S ratio 0.24 (Fig S1, right panel) have an average TM-score of only 0.59. To further verify this, each complex TM-score has been compared with the worst interface predictions derived from BIPSPI (Fig.S2, left panel). This comparison displayed a spearman correlation coefficient of 0.48 between training set similarity and interface prediction performance. Therefore, the excellent performance of BIPSPI is at least partially a result of structural similarity between parts of its training set and our test set. However, even considering low similarity complexes only (TM-score < 0.6) BIPSPI still yields the best performance between all the considered predictors (Fig.S2, right panel), i.e. overfitting is not the only factor causing this predictor superiority.

### Docking with the constraints from the binding site predictions

Next, we examined the ability to use the interface predictions to score docking models. Docking models from the GRAMM scan stage (GRAMM baseline) were re-scored using the interface probabilities (Eq. 1) from the interface predictors listed in Table 1. For comparison, we have also considered docking models rescored by the AACE18 potential^37^. A summary of the results is shown in Figure 5 and Supplemental Table S1. The most near-native docking models are top-ranked using the BIPSPI predictions, reaching SR(10) ~ 25% and SR(1) ~ 13%. Rescoring with this interface predictor is better than using the AACE18 potential (SR(10) ~ 18 % and SR(1) ~ 7 %).

**Figure 5.**
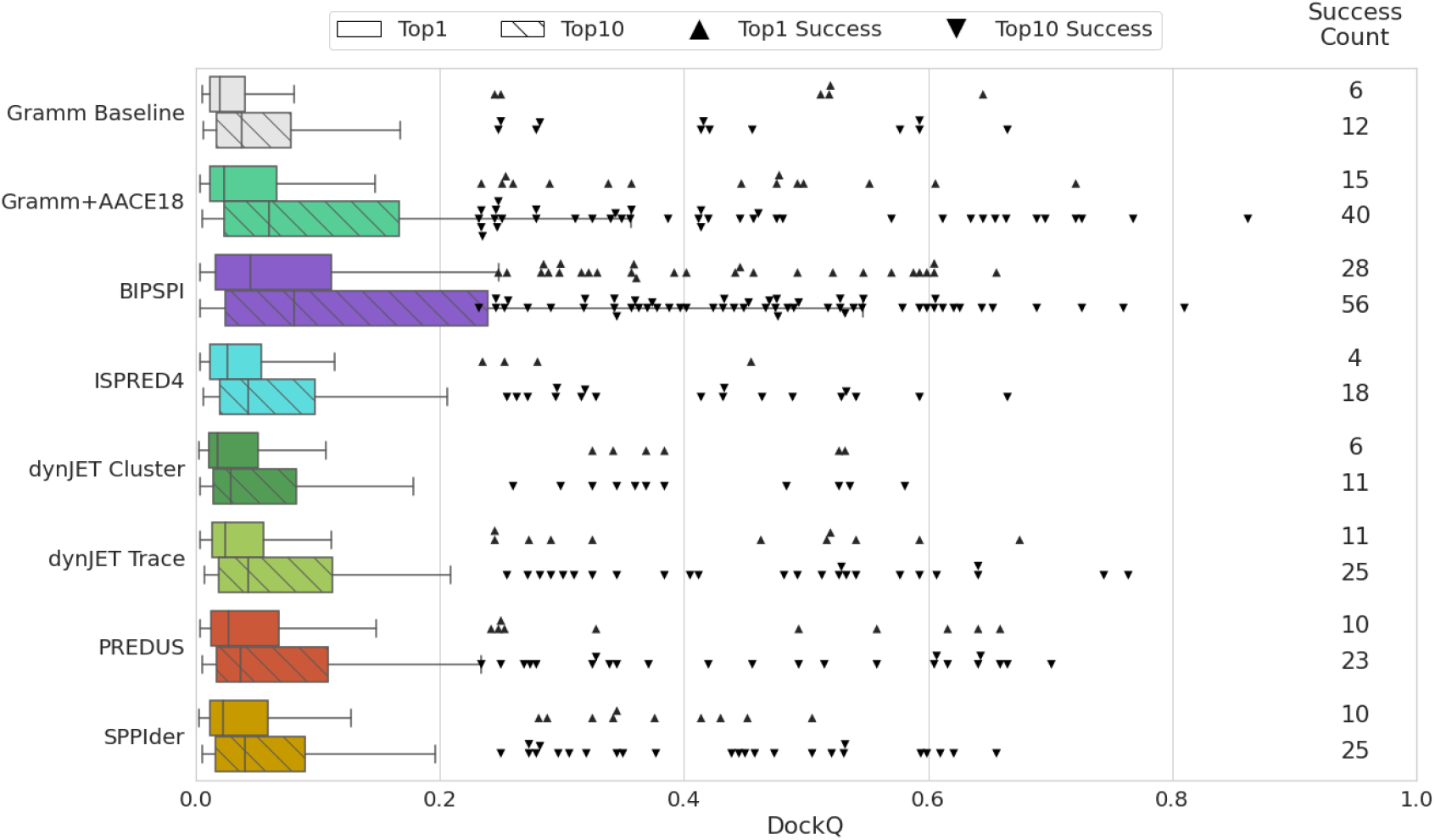
Performance of docking with constraints obtained by different interface predictors on 220 binary complexes from DOCKGROUND benchmark set 4. Horizontal bars represent the box-and-whiskers distributions of the DockQ scores, and each point represents a successful docking model (DockQ > 0.23). Non-striped bars and upward-pointing triangles display results obtained for the top-ranked docking models, while striped bars and downward-pointing triangles pertain to the model with the best dockQ score among the top 10 docking models. Pairs of success counts represent the number of targets for which successful docking models were generated in the corresponding docking run within top 1 (upper number) and top 10 (lower number) docking models.

Re-scoring the docking models with predictions from the other predictors does not significantly improve overall docking performance compared to the docking with scoring by shape complementarity only (GRAMM baseline in Fig 5), and they are far from the performance level of the AACE18. However, those predictors identify different top-ranked models than GRAMM. A partial overlap between the rescored list and GRAMM baseline docking is observed among the top 10 docking models (four cases for the ISPRED4 and one for the dynJET cluster). DynJET Trace, SPPIDER and PREDUS predictions yield a slight improvement over the baseline docking. Comparative analysis of predictor-driven docking reveals that different predictors, as a rule, move up near-native docking models for different complexes. Comparing interface predictors-based scorings (Fig. S3), only in one case (PDB 2bwe) 5 out of 6 predictor-driven dockings brought an acceptable model to the top of the prediction list. Further, top-1 acceptable docking models were obtained by four predictors only for three other complexes (PDBs 1b27, 1nbf and 1yu6). Thus, although the general impact of most predictors is low, there is a certain degree of complementarity between them, and their joint utilisation could enhance cumulative docking success significantly.

There are 15 complexes for which BIPSPI constraints failed to produce a top-1 near-native docking model while other interface predictors succeeded. Two of the complexes exhibit dockQ score < 0.02 for the top-ranked BIPSPI dockings (PDBs 3k9m, 3lwn). These “extreme” failures, together with one additional case (PDB 4pj2), are caused by a failed interface prediction of BIPSPI. For all other cases, the BIPSPI overall interface prediction quality is comparable to the best other method or better. Thus, failures here seem to be caused by BIPSPI tendency to be very precise (high PPV) at the expense of prediction completeness (data not shown). This leads to the number of generated (although correct) interface constraints being too weak to avoid significant rotational freedom between the two interacting patches. Note that considering acceptable models from top-10 docking models did not increase consistently the number of complexes for which constraints from the most predictors lead to the successful docking.

Finally, it should be noted that BIPSPI and AACE18 scoring are complementary to each other. Only four near-native top-1 complexes are shared, while BIPSPI and AACE18 separately succeeded for another 24 and 11 complexes, respectively (Fig S3F). When considering near-native docking models from the top 10 docking solutions, that overlap is slightly more considerable (19 common cases compared with 40 unique for AACE18 and 56 for BIPSPI).

### Simulated predictions

Various algorithms tested in this study produce interface predictions with TPR and PPV varying from protein to protein. Thus, in order to test the performance of the docking protocol in a controlled scenario (i.e. at pre-defined TPR and PPV values, which are the same for all complexes in the dataset), we introduce certain levels of “noise” into the native interface (see Methods). We have added noise by reducing PPV, i.e. adding false interface residue, and reducing TPR, i.e. removing correct interface residues. Results are reported in Figure 6 (top panel) and Supplemental Table S3.

**Figure 6.**
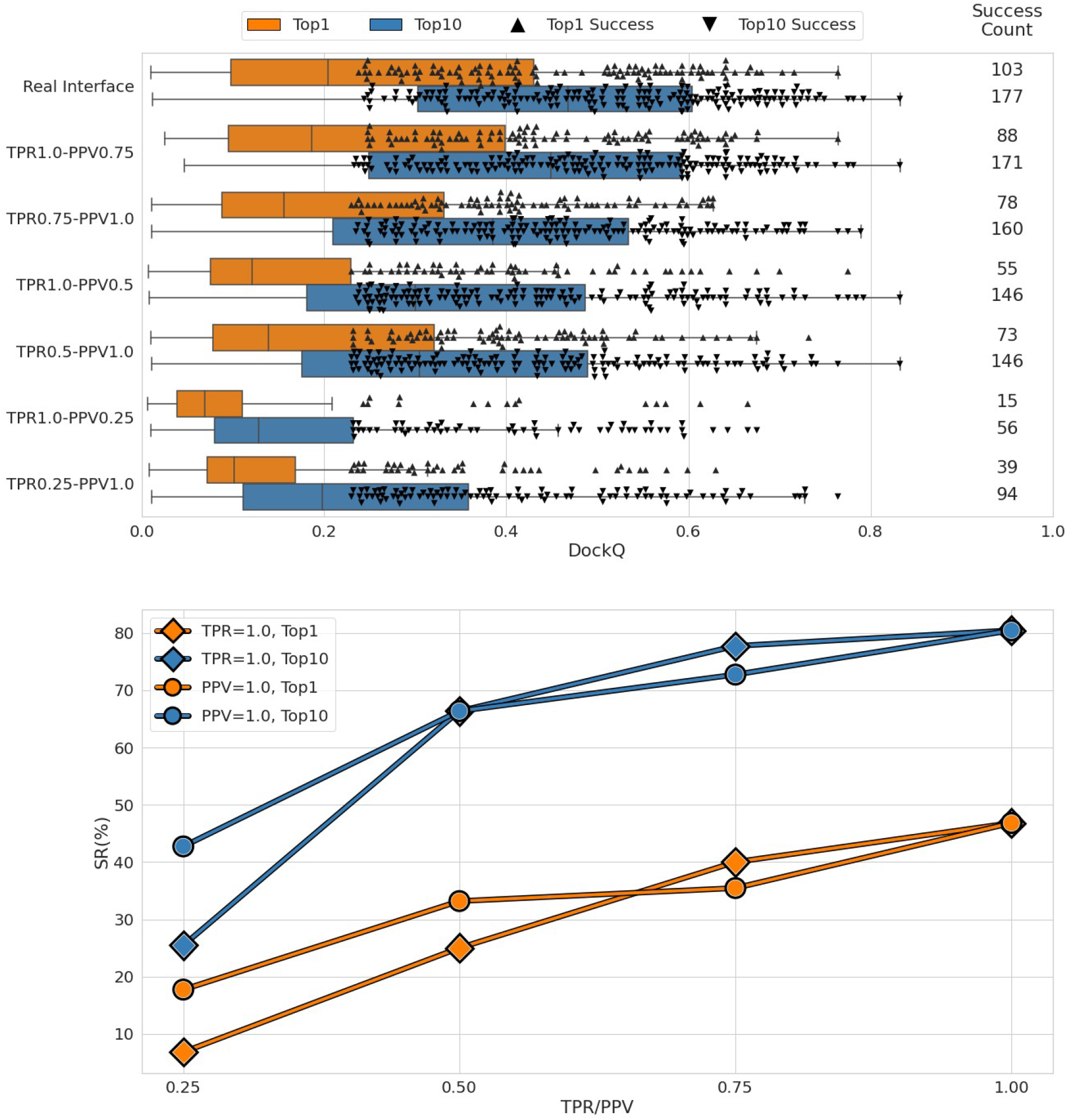
Performance of docking with constraints obtained from different simulated interfaces on 220 binary complexes from DOCKGROUND benchmark set 4. In the top panel, horizontal bars represent the box-and-whiskers distributions of the DockQ scores, and each point represents a successful docking model (DockQ > 0.23). Orange bars and upward-pointing triangles display results obtained for the top-ranked docking models, while blue bars and downward-pointing triangles pertain to the model with the best dockQ score among the top 10 docking models. Pairs of success counts represent the number of targets for which successful docking models were generated in the corresponding docking run within top 1 (upper number) and top 10 (lower number) docking models. The bottom panel displays success rates for top 1 (orange) and top 10 (blue) docking models obtained for a series of simulated interfaces with varying PPV (diamonds) or TRP (circles) while another parameter (TPR or PPV, respectively) is kept 1. Lines are guides for the eye.

In general, docking success is reduced by both under- (false negatives) or over- (false positives) interface predictions. In the scoring scheme used in the paper (Eq. 1), the contribution of a large patch of true interface residues (covering the entire interface, TPR = 1) overweights the contribution from a small amount of wrongly predicted non-interface residues (PPV=0.75). On the other hand, even relatively small under-prediction of the interface (TRP=0.75) gives rise to the undesired energetical ‘freedom’ in the ligand placement even in the absence of wrongly predicted non-interface residues (PPV=1). The trend is reversed when the level of ‘noise’ at the predicted interfaces increases, and this behaviour is the same for both top 1 and top 10 docking models.

### Complex-wise analysis

The DockQ score for the docking models exhibits a strong correlation to the AUC of the interface predictors for the corresponding protein chains. (Fig.7). Few exceptions are found in complexes with high shape complementarity (Fig.8A), which is sufficient in some cases to achieve acceptable dockings even with low-quality constraints. Another possibility to obtain good dockings from noisy constraints is the combination of wide scattering of false positives predictions over the entire surface and tightly packed true positives. Such scattering, observed, for instance, in dynJET2 Trace predictions, allows in some cases successful docking regardless of somewhat inaccurate predictions (data for PDB 3bx1 in Table S3).

**Figure 7.**
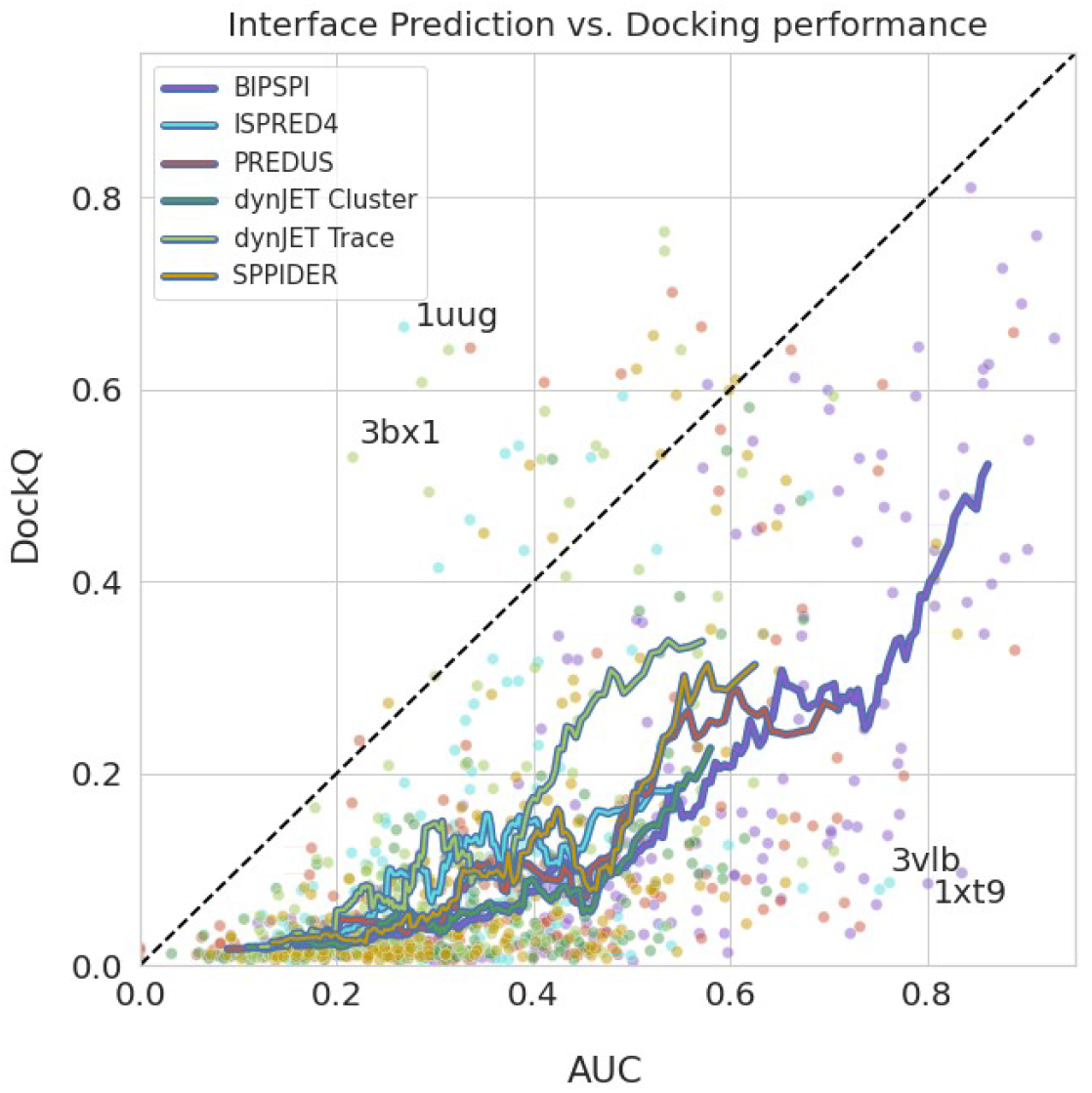
Correlation between area under Precision-Recall curve (AUC) and the DockQ score of the best among top 10 docking models for various interface predictors. Data are shown only for those 177 binary complexes from DOCKGROUND benchmark set 4 with near-native docking models in the upper baseline docking (constraints derived from the native interfaces). For each complex, averaged receptor and ligand’s AUC are plotted. Running averages have been obtained from a sliding window of 20 data points.

**Figure 8.**
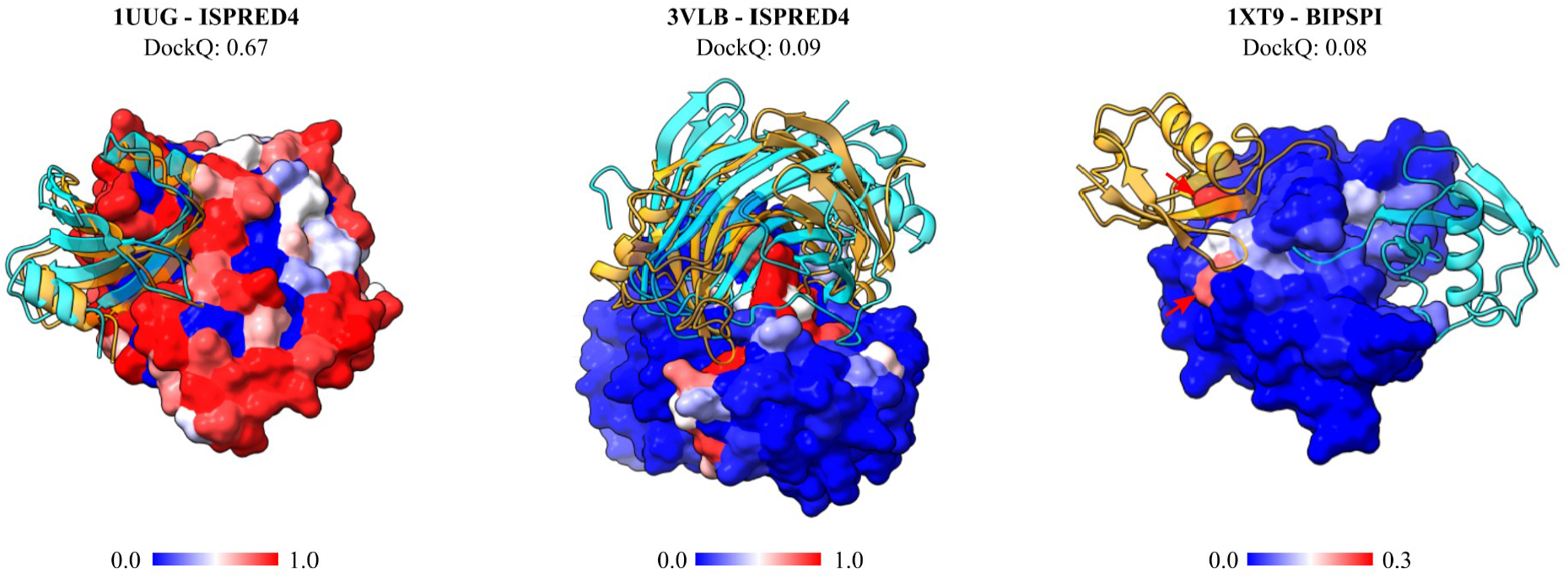
Three examples of constrained docking results; receptor is represented by atomic surfaces; all receptor atoms are colored according to the score given from the interface predictor, with blue to red transition indicating low to high scores; two ligands depicted in cartoon style are present in each structure, one in the native position colored in cyan and one resulting from the docking procedure (best in top 10 ranks), colored in orange; (left) acceptable docking compared to the bound structure with PDB ID 1uug yielding a DockQ score of 0.67, resulted from application of ISPRED4 scorings as constraints; (middle) example of incorrect docking due to rotational ambiguity in the relative orientation of the correct interfaces; docking produced using ISPRED4 scores (which yields a DockQ score of 0.09) is rotated of 180° around a vertical axis passing through the center of the receptor, respective to the native bound structure with PDB ID 3vlb; (right) example of incorrect docking due to high scoring of residues close but not belonging to the actual interface; the docking (with DockQ score of 0.02) is obtained adopting BIPSPI predictions and compared to the bound structure with PDB ID 1xt9; nevertheless the average 0.8 AUC value for this complex interface predictions, False Positive residues (indicated with red arrows) with high scoring are enough to drive the docking toward a completely wrong configuration. Those residues are part of the receptor catalytic site.

Constraint quality in our protocol also seems to be an essential but not sufficient condition for successful docking. A significant number of complexes exhibit low DockQ scores (~0.1) for the best out of the top 10 docking models, even with large AUC values (Fig 7). In those docking models, the ligand is placed into or close to the correct binding site of the receptor with the correct patches of ligand and receptor residues facing each other but with a wrong mutual orientation (Fig.8B). Subsequently, rotational freedom is quite a common pitfall of using interface constraints predicted independently for the receptor and ligand and can be seen as an intrinsic limitation of this method.

A fascinating case is given by the complex between the Den1 protease and Nedd8, a Ubiquitin-like protein (Fig.8C). The biological role of this complex is to activate Nedd8 by removing a portion of its disordered C-terminal ^44^. Predicting the interface of this complex with BIPSPI identifies a strong signal in the protease catalytic triad residues, shown by arrows in Fig 8C. These residues are located at the very edge of the interface and have better scores than the other predictions in the interface of Den1. The highest scores for the Nedd8 predictions are obtained for the amino acids in the middle of the Nedd8 interface. Since the peripheral of the Den 1 interface is located far away from the central part of its interface, docking poses with those high-scored predictions facing each other and thus favoured by the scoring scheme (Eq 1) are incorrect with the location of the ligand far away from its native position (Fig 8C).

## CONCLUSIONS

In this work, we analysed the use of predicted interface residues for scoring template free docking solutions. First, we show that interface information is sufficient to correctly identify an acceptable model for the vast majority of all targets that could be generated. Using predicted interfaces, we found that one predictor, BIPSPI, clearly was superior to all the other predictors. Using the interfaces predicted from BIPSPI, almost twice (12.5% vs 6.8%) as many first ranked models are acceptable (dockQ>0.23). In contrast to the other predictors, BIPSPI uses two chains to predict interface contacts and predicts the probability for each pair of residues to be in contact. In conclusion, a general framework to test interface prediction as a constraint for protein docking has been produced in this study. This framework has shown a promising potential on a large number of docking cases and, thanks to its simplicity and flexibility, it may be easily adapted to use any kind of interface prediction, hopefully helping improve the state of the art of protein-protein docking.

## ACKNOWLEDGEMENTS

We thank the Swedish National Infrastructure for Computing for providing computational resources. This work was supported by a grant VR-NT-2016-03798 from the Swedish National Research Council (www.vr.se) to AE. The salary of PK, and GS were partly paid by grants from the Swedish Research Council.

## SUPPLEMENTARY MATERIAL

**Table S1.** Summary of the docking results with constraints derived from the various interface predictors.

| Docking | Top 1 |  |  |  | Top 10 |  |  |  |
| --- | --- | --- | --- | --- | --- | --- | --- | --- |
|  | Median | Mean | St.Dev | SR(1),% | Median | Mean | St.Dev | SR(10), % |
| Gramm Baseline | 0.02 | 0.04 | 0.08 | 2.7 | 0.04 | 0.07 | 0.10 | 5.5 |
| Gramm + AACE18 | 0.02 | 0.07 | 0.11 | 6.8 | 0.06 | 0.13 | 0.17 | 18.2 |
| BIPSPI | 0.04 | 0.10 | 0.14 | 12.7 | 0.08 | 0.17 | 0.20 | 25.5 |
| ISPRED4 | 0.03 | 0.04 | 0.05 | 1.8 | 0.04 | 0.09 | 0.12 | 8.2 |
| PREDUS | 0.03 | 0.06 | 0.10 | 4.6 | 0.04 | 0.10 | 0.14 | 10.5 |
| dynJET Cluster | 0.02 | 0.04 | 0.07 | 2.7 | 0.03 | 0.07 | 0.09 | 5.0 |
| dynJET Trace | 0.02 | 0.06 | 0.10 | 5.0 | 0.04 | 0.10 | 0.15 | 11.4 |
| SPPIDER | 0.02 | 0.05 | 0.08 | 4.5 | 0.04 | 0.09 | 0.14 | 11.4 |

**Table S2.**
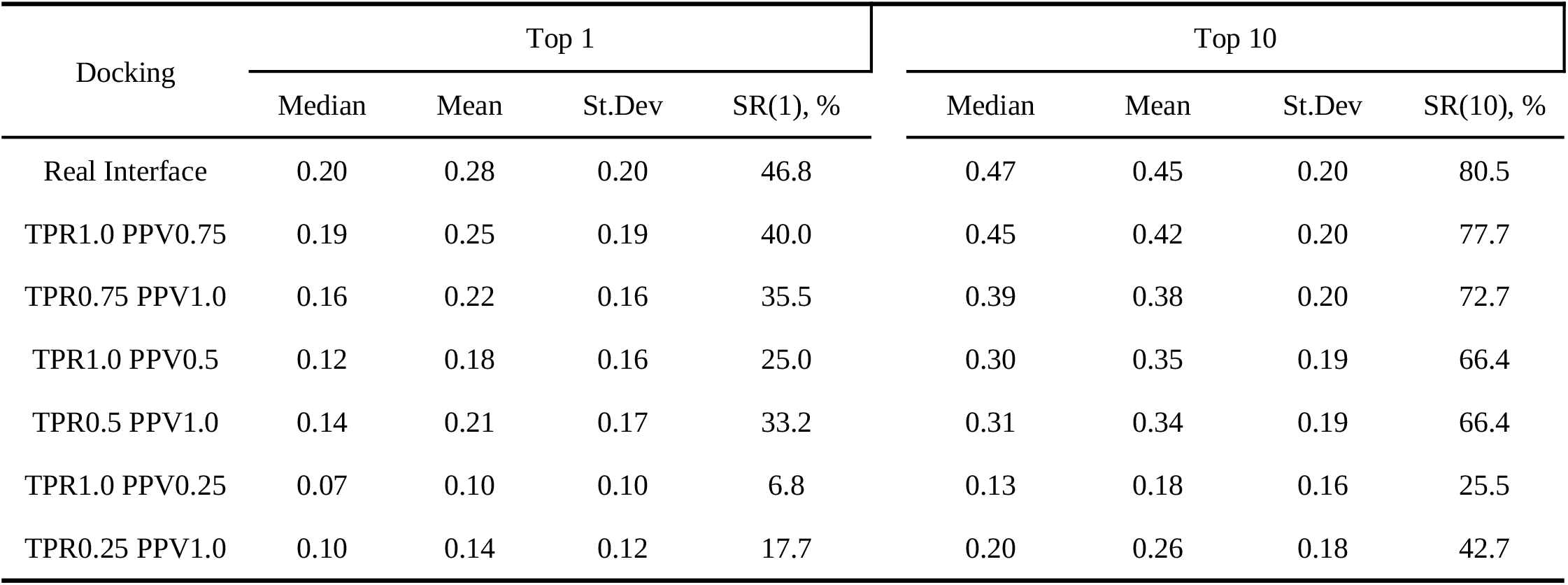
Summary of the docking results with constraints derived from the simulated interfaces.

**Figure S1.**
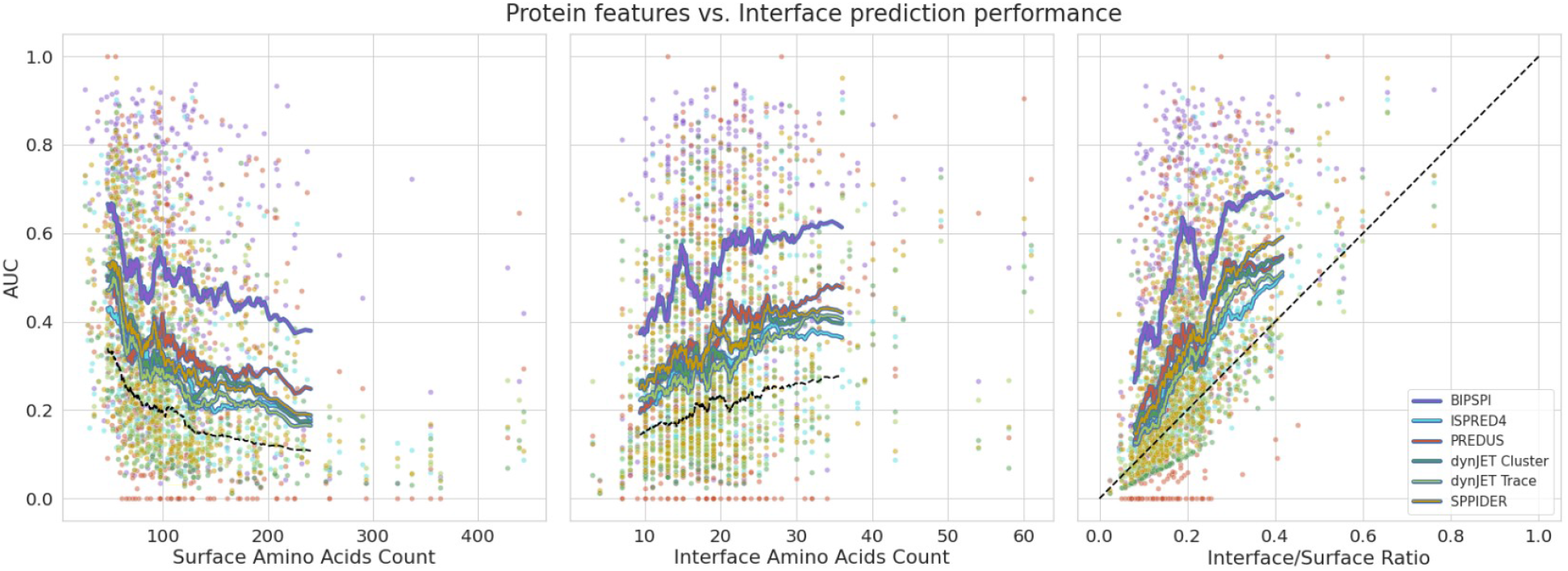
Impact of single proteins features over interface prediction performance. Each point represents a protein from the DOCKGROUND dataset, colored accordingly to the predictor adopted to obtain interface estimates. Running averages have been calculated in both cases considering a window of 50 proteins sliding along the independent variable axis. Dashed lines indicate the expected random predictor performance in terms of interface to surface residues ratio. In the left panel, the number of surface residues in single proteins (determined by relative solvent accessibility value > 0.2) has been plotted against the related interface prediction AUC for all considered predictors. In the middle panel, the number of residues which belong to the native complex interface has been plotted instead against AUC values. Finally, in the right panel, the ratio between the number of surface residues and the number of residues which are part of the interface has been plotted as well, against the interface prediction AUC.

**Figure S2.**
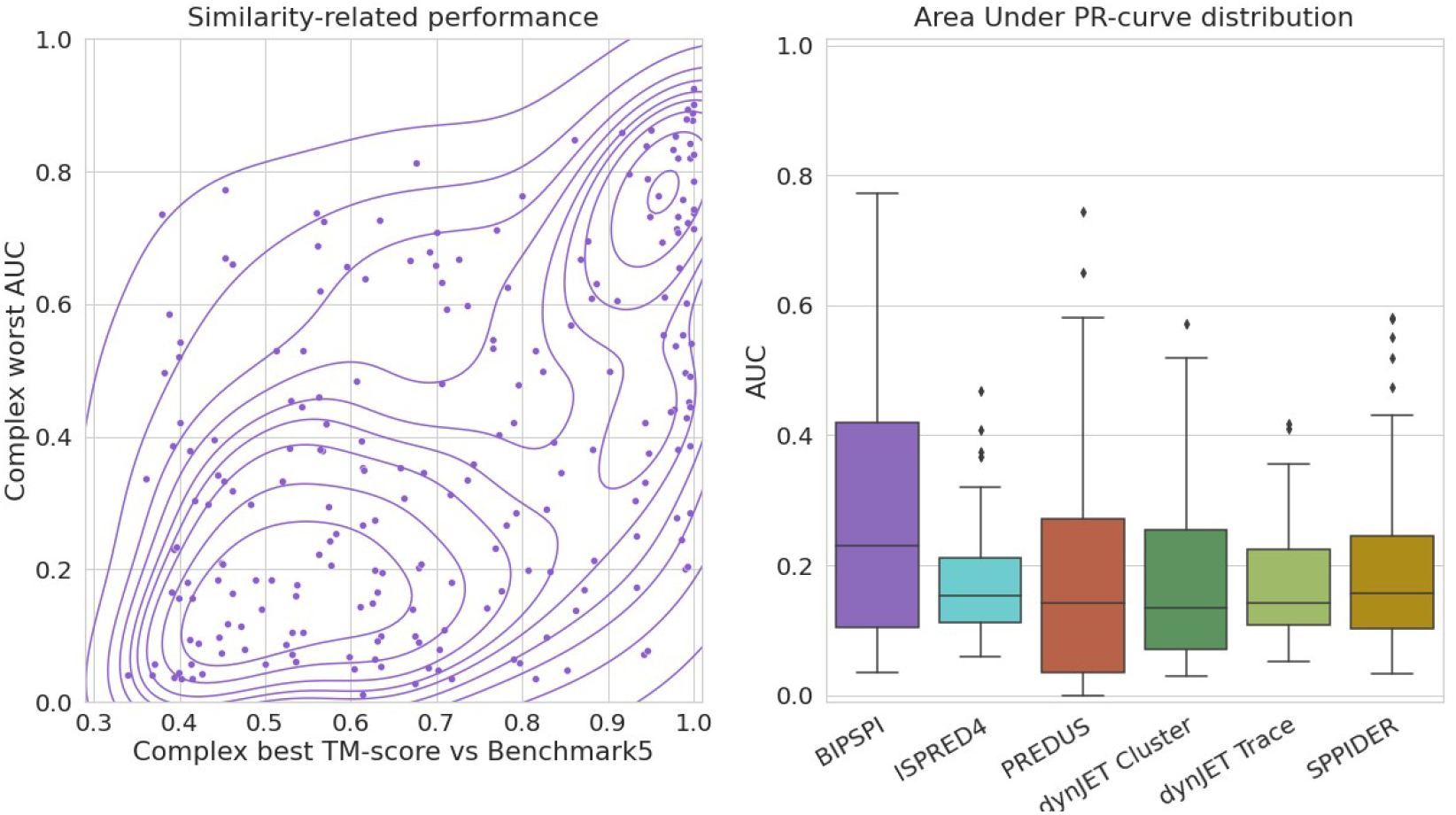
Correlation of interface prediction performance to BIPSPI training set similarity. Left panel shows the correlation between the highest TM-score obtained comparing DOCKGROUND and Benchmark5 complexes and the worst chain predictions generated by BIPSPI for the same complexes. Kernel density estimates lines provide a visual help to define point density. The right panel reports distributions, for each predictor, of each complex worst chain prediction, for each complex with TM-score against Benchmark5 lower than 0.6.

**Figure S3.**
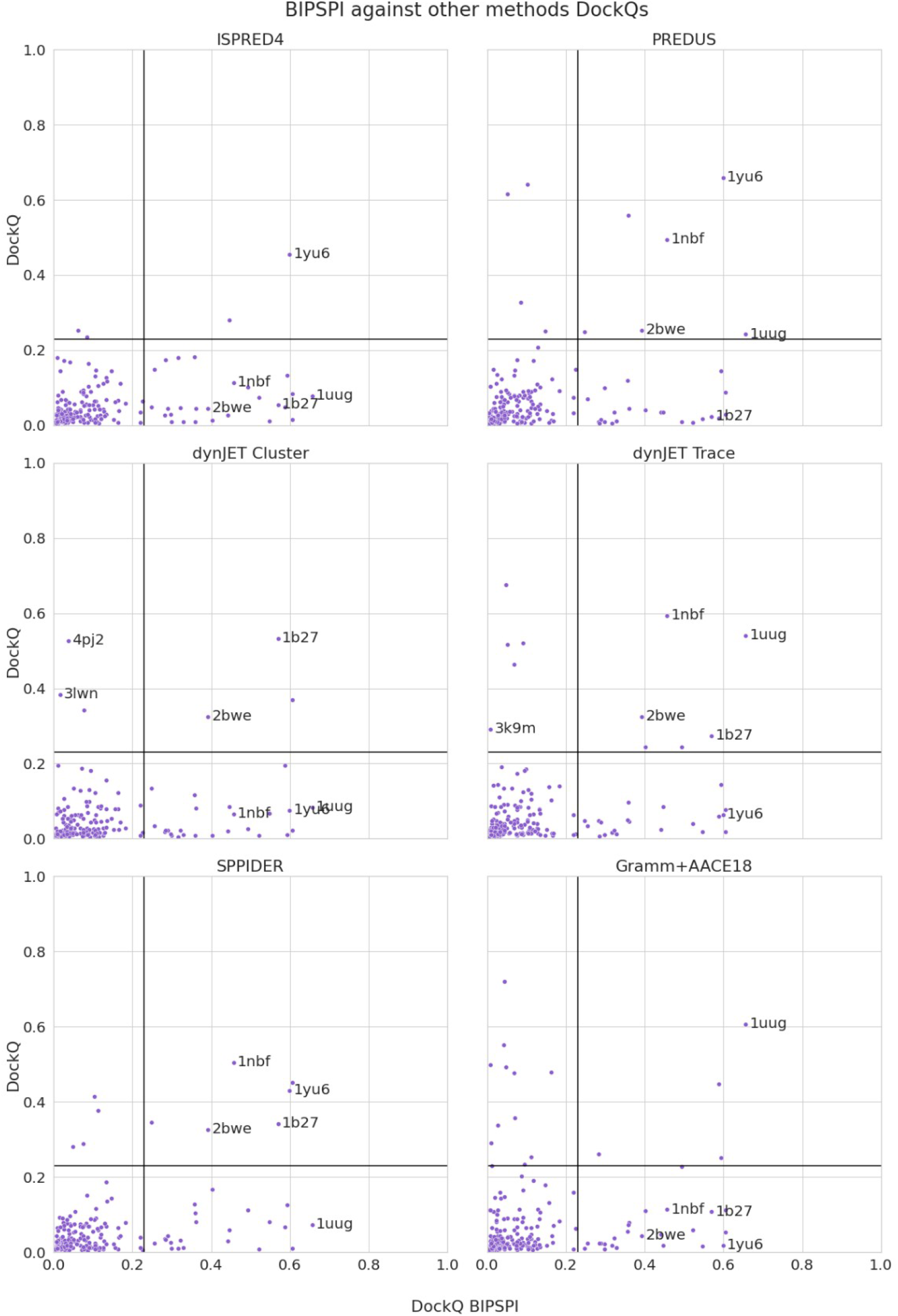
Correlation of BIPSPI and other predictors top ranked docking results. Scatterplot showing for each complex the top ranked docking DockQ scores obtained using BIPSPI constraints, versus the scores obtained using other predictors constraints. Particular docking cases with acceptable quality (DockQ score >= 0.23) with at least one predictor have been annotated with the respective PDB ID. Horizontal and vertical lines indicate the thresholds to consider a docking acceptable.

**Table S3.**
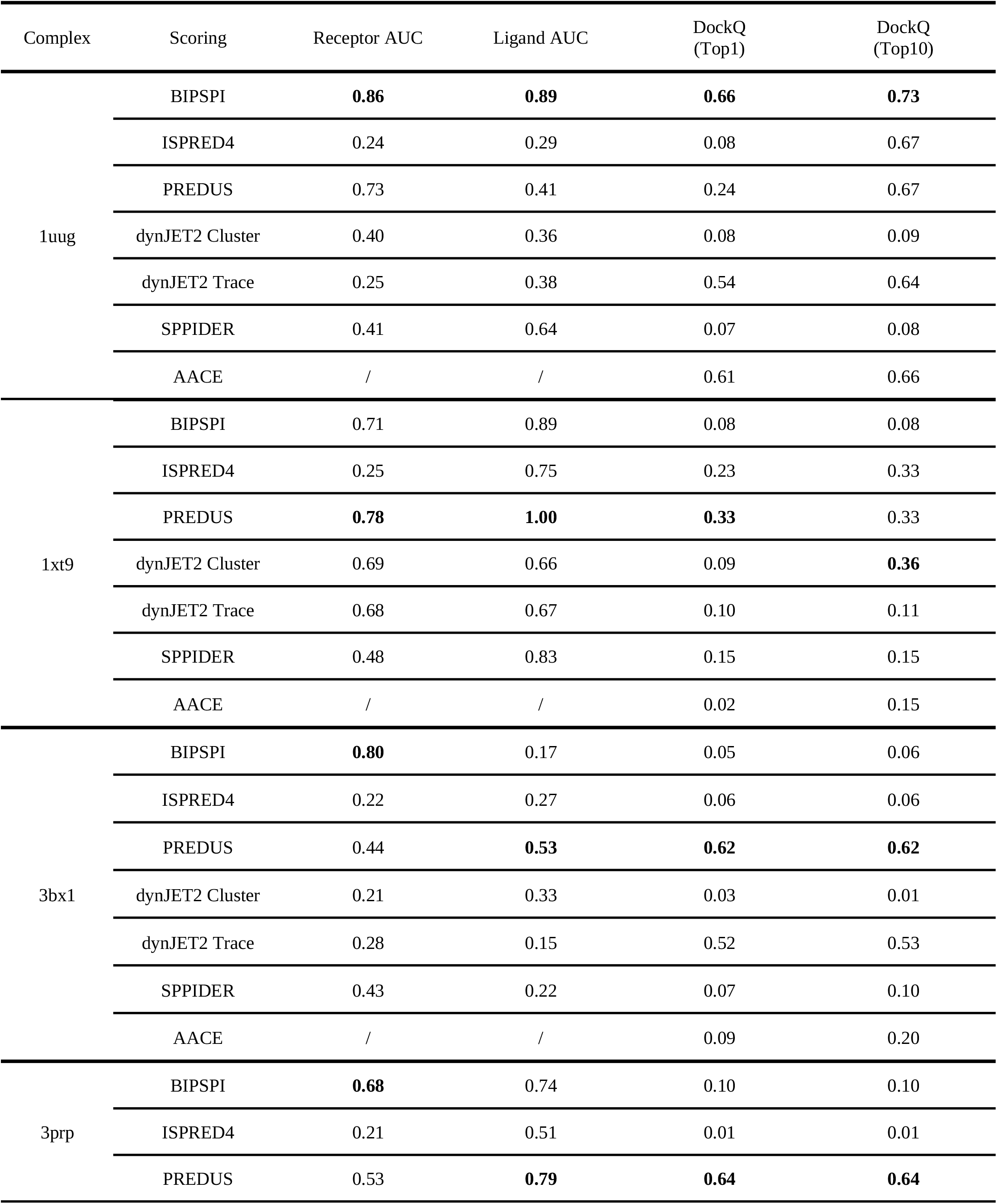
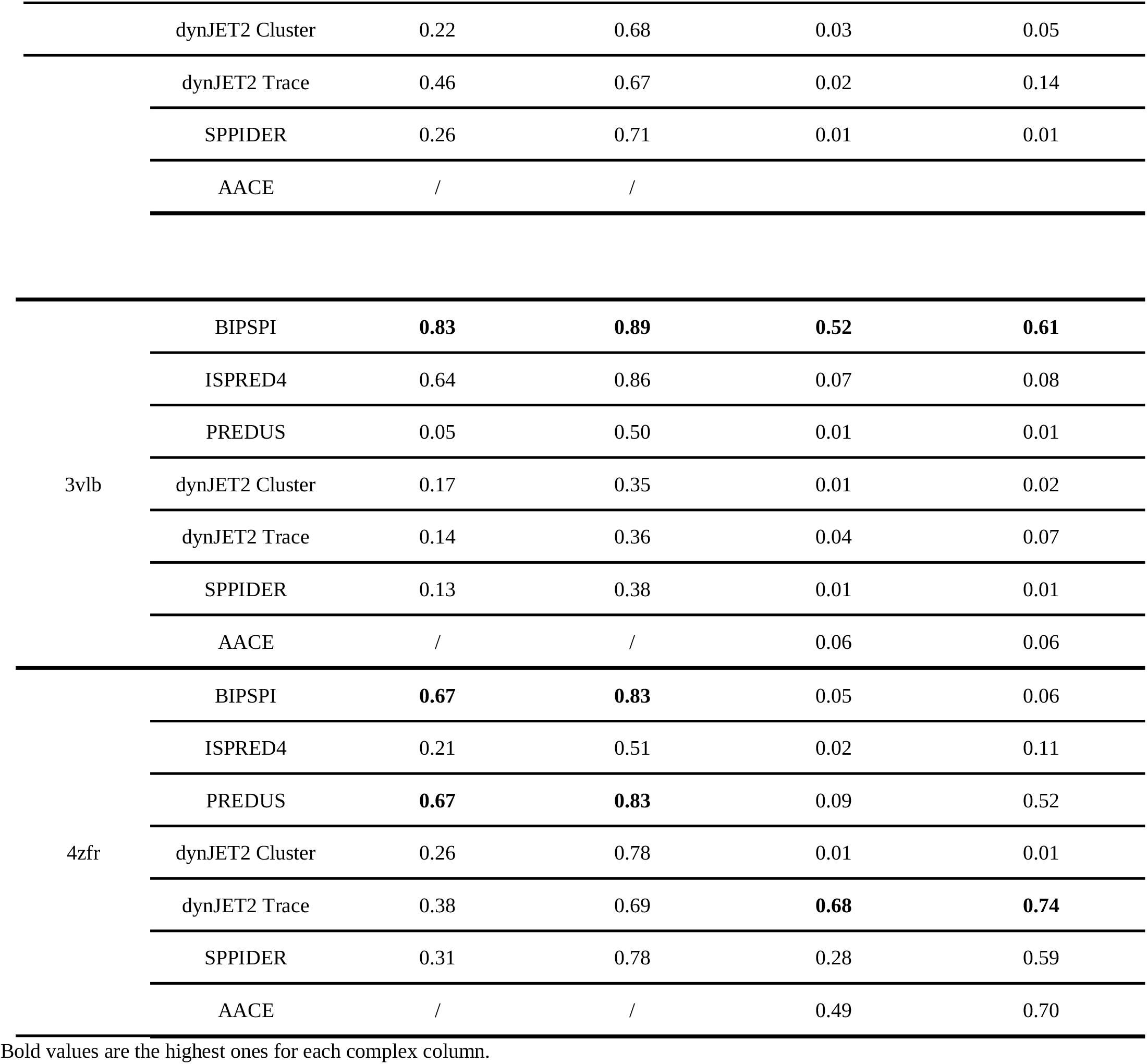
Summary of the docking results and predictions AUC for individual complexes

## Notes

### Competing Interest Statement

The authors have declared no competing interest.

